# Variation in context dependent foraging behavior across pollinators

**DOI:** 10.1101/210377

**Authors:** Heather M. Briggs, Stuart Graham, Callin M. Switzer, Robin Hopkins

## Abstract

Pollinator foraging behavior has direct consequences for plant reproduction and has been implicated in driving floral trait evolution. Exploring the degree to which pollinators exhibit flexibility in foraging behavior will add to a mechanistic understanding of how pollinators can impact selection on plant traits. Although plants have evolved suites of floral traits to attract pollinators, flower color is a particularly important aspect of the floral display. Some pollinators show strong innate color preference, but many pollinators display flexibility in preference due to learning associations between rewards and color, or due to variable perception of color in different environments or plant communities. This study examines the flexibility in flower color preference of two groups of native butterfly pollinators under natural field conditions. Our study reveals that pipevine swallowtails and skippers, the predominate pollinators of the two native Texas Phlox species, display distinct patterns of color preferences across different contexts. Pipevine swallowtails exhibit highly flexible color preferences and likely utilize other floral traits to make foraging decisions. In contrast, skippers have consistent color preferences and likely use flower color as a primary cue for foraging. As a result of this variation in color preference flexibility, the two pollinator groups impose concordant selection on flower color in some contexts but discordant selection in other contexts. This variability could have profound implications for how flower traits respond to pollinator-mediated selection. Our findings suggest that studying dynamics of behavior in natural field conditions is important for understanding plant-pollinator interactions.

## Introduction

Plants solicit pollinator visitation through a variety of floral display traits including color, scent, size, and shape (Schiestl and Johnson 2013). These traits generally advertise to pollinators the availability of a reward such as pollen and nectar. Extensive studies, both in controlled laboratory settings and natural field conditions, have documented pollinators disproportionately visiting flowers with particular trait values (Schemske and Bradshaw 1999; Byers et al. 2014; Thairu and Brunet 2015). Pollinator foraging choices can result from innate sensory sensitivity to a particular trait state; but this innate preference can be flexible due to a number of factors including learned associations between particular trait states and a reward (Weiss 1997; Gumbert 2000; Goulson et al. 2007; Raine and Chittka 2007a). Most studies investigate pollinator visitation patterns in reference to variation in a single floral trait in a single environment, and yet there is evidence that interactions between traits and environmental context can alter pollinator preference (Hersch and Roy 2007; Raguso 2008; Pohl et al. 2011; Yoshida et al. 2015). Despite the numerous factors that contribute to pollinator behavior, we know surprisingly little about the degree to which pollinators are consistent or flexible in their preference for a specific trait across natural variation in floral displays and under natural field conditions.

Flower color is a particularly important trait driving patterns of pollinator visits to plants (Frisch et al. 1914; Ômura and Honda 2005; Dötterl et al. 2014). The relationship between pollinator identity and preferred flower color is so widely observed that it is often assumed that flowers of a particular color are visited by specific pollinators (e.g. red flowers are pollinated by hummingbirds and white flowers are pollinated by moths or bats). Extensive pollinator experiments both in natural field conditions and in controlled laboratory conditions show some support for broad patterns of general color preference for particular pollinators (Waser et al. 1996; Muchhala 2003; Fenster et al. 2015). For example some studies show that: bees tend to prefer blue or yellow flowers (Giurfa et al. 1995), butterflies often prefer blue or yellow (Weiss 1997), and hummingbirds tend to visit red flowers (Shrestha et al. 2013). Much of these associations between flower color and pollinators are thought to be due to the specificity of visual systems in pollinator taxa. Birds, mammals, and insects differ in their visual abilities across the UV and visible color spectrum and this variation can explain some of the innate differences in color preferences across pollinator taxa (Briscoe and Chittka 2001). In other cases the external environment is also important for explaining patterns of pollinator color preference. For example, crepuscular and nocturnal nectar foragers such as hawkmoths and bats likely prefer white flowers because they are most visible under low light conditions (Johnsen et al. 2006). Despite these generalizations, there are many exceptions to these foraging patterns (Smith et al. 2008) and some pollinators appear to exhibit flexibility in color preference (Waser et al. 1996; Ollerton et al. 2009; Rosas-Guerrero et al. 2014). Understanding if and how pollinators show variation in their color preference across plant species and in different environments is important for understanding the eco-evolutionary dynamics of plants and pollinators in a community context.

Flexibility in flower color preference can be driven by a number of factors including learned associations between color and rewards, variation in environmental or community context, and interactions with other floral trait signals. Controlled laboratory studies indicate many pollinators including bees, flies and butterflies can alter their innate color preference by learning an association between nectar or pollen reward and a new color (Weiss 1997; Hollis and Guillette 2011; Blackiston et al. 2011). For example, while bumble bees often display an innate preference for blue flowers they can readily learn to associate a reward with a novel flower color (Raine and Chittka 2007b; Ings and Chittka 2009). Although rarely studied in the field, these lab studies suggest that learning could explain some of the observed variation in flower color preference we see in nature. Plant community characteristics can also contribute to variation in color preference. For example, variation in the degree to which flower color contrasts with the background can drive flexibility in color preference (Osorio and Vorobyev 2008). Additionally, some community factors such as presence of other flowering species or presence of a dominant pollinator competitor can contribute to context dependent flower color preferences (Brosi and Briggs 2013, Fornoff et al. 2016). Finally, some studies have shown that innate preference and learning of a particular trait, such as color, can vary depending on other aspects of the complex floral display such as scent, size and shape. For example, strength and direction of preference for a certain flower color can depend on presence or absence of scent signals (Leonard and Masek 2014; Knauer and Schiestl 2014; Yoshida et al. 2015; Russell et al. 2016). For these reasons, it is likely that pollinators in nature could display extensive flexibility in floral color preference across plant species and community contexts. Yet, the extent to which color preference is flexible versus consistent in nature is poorly understood.

The flexibility of pollinator color preference can have important implications for the evolution of floral traits in natural communities. It is well documented that pollinator preference leads to increased floral visitation and thus selection for the preferred flower type (Schemske and Bradshaw 1999; Aldridge and Campbell 2007). Despite the recognized importance of pollinator preference on plant trait evolution, we have little information regarding the consistency of these pollinator behaviors under field conditions. Often pollinator observations in the field are conducted on controlled arrays under limited contexts. Little attention has been given to how observed preference in one foraging context might vary in other contexts. Understanding flexibility in pollinator behavior can provide a deeper understanding of how traits evolve in plants to recruit effective pollination services. Much of our understanding of pollinator foraging behavior and subsequent color preferences come from studies focusing on bee pollinators. While bees are an important group of pollinators, particularly for agriculture, other insects such as flies and butterflies serve as essential pollinators in many natural communities. In general, we know much less about solitary foragers, and some studies suggest that what we know about the foraging behavior in social insects does not always apply to non-social pollinators (Kelber and Pfaff 1999). Therefore, studying other groups, such as Lepidoptera, can offer important insights into pollinator foraging behavior in non-social insects.

Lepidoptera are ideal for investigating flexibility in color preference because they have variation in visual systems, display innate color preference, and can alter preference through both learning and environmental context. In contrast to hymenopteran pollinators that exhibit little variation in visual photoreceptors, lepidopterans display substantial variation in photoreceptor spectral sensitivities (Briscoe and Chittka 2001). This has led to marked differences in both innate and learned color preferences across species of butterflies (Stavenga and Arikawa 2006; Briscoe 2008). Innate preferences undoubtedly play an important role in foraging decisions, but controlled laboratory studies show that some butterflies can be flexible with their color preference to improve their foraging success through learning color-reward associations (Kandori et al. 2009; Pohl et al. 2011; Blackiston et al. 2011). While largely unexplored, there is also some evidence that butterflies can have flexible color preference depending on the environmental context of the display. For example, male and female Papilo xuthus have disparate color preferences (purple and light blue respectively) when tested in a neutral background but when tested against a green background, both sexes prefer more reddish colors (Kinoshita et al. 1999). Despite the evidence that butterflies can be flexible in their color preference, very few studies have explored the extent to which they are flexible in their color preference in natural systems.

This study investigates the flexibility of flower color preference in two butterfly pollinators on two plant species in a natural field environment. The pipevine swallowtail (Battus philenor; hereafter pipevine swallowtails) and a variety of skipper species (family Hesperiidae; hereafter skippers) are the primary pollinators of two Texas wildflower sister species: Phlox drummondii and Phlox cuspidata (Hopkins and Rausher 2012, Hopkins and Rausher 2014). Throughout its range, P. cuspidata has light-blue flowers characteristic of most Phlox species. P. drummondii also has the same light-blue flower color across much of its range, but in some eastern and central Texas populations P. drummondii has evolved dark-red, light-red, and dark-blue flower colors (Hopkins and Rausher 2011). Pipevine swallowtails and skippers make up around 95% of the total visits to the two plant species, and have been shown to visit all of the P. drummondii flower colors. As such, this system provides an opportunity to decouple flower color from plant species identity and thus explore how flower species context influences the flexibility of color preference in two groups of butterfly pollinators.

We observed the foraging behavior of both pipevine swallowtails and skippers on experimental arrays composed of combinations of the two Phlox species and varying flower colors morphs of P. drummondii in natural field settings. Specifically, we explore the flexibility in color preference across three plant foraging contexts: (i) within species color variation (when two P. drummondii plants differ in color) (ii) different species color variation (when P. drummondii and P. cuspidata differ in color) (iii) within species color variation with other species present (when two P. drummondii plants differ in color but P. cuspidata is also present in the array). With these arrays we address two specific questions: (1) Is pollinator color preference flexible depending on foraging context (i.e. species context or the presence of an additional species with similar phenotype)? (2) Do the two primary pollinators of Phlox show similar color preference across different contexts? The power of our experiment comes from answering these questions across three different color comparisons.

## Material and Methods

### Study System

Phlox is a butterfly-pollinated genus (Levin and Berube 1972). P. drummondi and P. cuspidata are annual herbs native to central and eastern Texas that inhabit roadsides, open fields, and pastures. Individuals germinate in late fall or early spring and flower and set fruit from March through June. Both P. cuspidata and P. drummondii are predominantly pollinated by pipevine swallowtails and a variety of skipper species [family Hesperiidae; (Hopkins and Rausher 2012)].

P. drummondii has four flower color variants in nature: light-blue, dark-red, light-red, and dark-blue. Butterfly foraging behavior generates selection on flower color and maintains the flower color polymorphisms across the P. drummondii range. In western populations, P. drummondii individuals have light-blue flowers; however, in populations sympatric with the light-blue-flowered P. cuspidata, P. drummondii has evolved dark-red flowers (Hopkins and Rausher 2011; Hopkins and Rausher 2012). In a small geographic region where light-blue and dark-red P. drummondii meet, all four flower colors can be found (Hopkins et al. 2014). P. drummondii individuals with different flower colors do not differ systematically in other traits such as nectar, scent, shape, or size [Hopkins, unpubl. data]. In sympatric populations, pollinator behavior causes greater reproductive isolation between darkred P. drummondii and P. cuspidata than light-blue P. drummondii and P. cuspidata, thereby favoring dark-red flower color through reinforcing selection (Hopkins and Rausher 2012). In the absence of P. cuspidata, pollinators prefer light-blue over dark-red flower color, thus selecting for light-blue flower color in allopatric populations (Hopkins and Rausher 2014).

The plants used in this experiment were collected from natural populations throughout the native ranges of P. drummondii and P. cuspidata. Plants were grown from seed in greenhouses at the Arnold Arboretum of Harvard University (2015) and at the University of Texas, Austin (2010 and 2012). To synchronize germination, we soaked seeds in 500 ppm gibberellic acid for 48 h, planted them in water-saturated Metro-Mix 360 (Sun Gro Horticulture, Bellevue, WA), and stratified them at 4 °C for 7 days. Plants were allowed to germinate and grow in growth chambers set for 14 h day-light and a 25 °C/22 °C day/night temperature regime. Plants were watered as needed and fertilized regularly with DynaGro Liquid Bloom fertilizer (Dyna-Gro Nutrition Solutions, Richmond, CA, USA). All plants considered to be healthy were transported to The University of Texas Brackenridge Field Laboratory (Austin, TX, USA) for experimentation.

### Pollinator observations

We assessed butterfly color preference over three years (2010, 2012, and 2015) in the month of May at the Brackenridge Field Laboratory (Austin, TX, USA). This site is located in the allopatric range of P. drummondii and wild populations of light-blue P. drummondii exist nearby. The pollinator observations from 2011 and 2012 are included in previous publications investigating selection on Phlox (Hopkins and Rausher 2012; Hopkins and Rausher 2014).

We recorded the foraging behavior of free-flying butterflies on arrays of live Phlox plants to examine color preference in varying contexts. We conducted pollinator observations on at least two days per array between 10 a.m. and 4 p.m. Each pollinator was identified as pipevine swallowtail, skipper, or ‘other’. While many skipper species have been recorded at the Brackenridge Field Laboratory we estimate, based on visual recognition, that only five of those species (Thorybes pylades, Erynnis horatius, Copaeodes aurantiaca, Atalopedes campestris, Hylephila phyleas) likely visited our arrays. Skipper butterflies are difficult to identify on the wing and all five species are of similar shape and size. For each pollinator we recorded the color and species of each flower visited. Pollinator visits were counted only if the pollinator’s proboscis was seen entering a corolla. From these data we calculated the total number of plants visited of each color by each pollinator.

#### Experimental Arrays

Each experimental array represented one of three plant community contexts (described below). Every array contained light-blue flowers of a focal species and an equal number of *P. drummondii* flowers of one other color (light-red, dark-red, or dark-blue). Across the arrays we varied both the species identity of the light-blue flowers and the color of the non-light-blue P. drummondii flowers in a full factorial design to give a total of nine distinct experimental array types (see Fig. 1). Our three foraging contexts were:

**Fig. 1.**
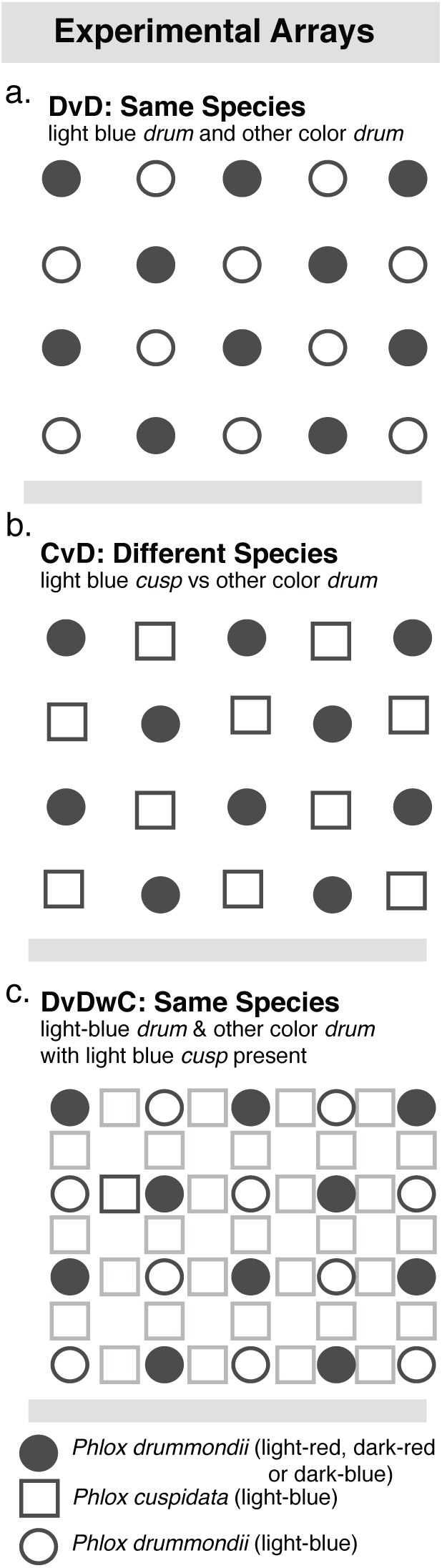
Schematic of the pollinator observation arrays placed in the field. For each array type we alternated focal flower colors in a 4 × 5 grid. Grey boxes (P. cuspidata) in panel c. indicate that plants were present, but pollinator foraging data were not collected on these plants. DvD data were collected in 2010, DvDwC, in 2012 and CvD in 2015.

##### (1)same species arrays

light-blue P. drummondii and other color P. drummondii, hereafter DvD (2) different species arrays: light-blue P. cuspidata and other color P. drummondii, hereafter DvC (3) same species arrays: light blue P. drummondii and other color P. drummondii with light blue P. cuspidata present in array, hereafter DvDwC.

Context (1) and (2) were the same except that the species of light-blue flowers differed. Context (1) and (3) were the same except for the presence/absence of P. cuspidata. Foraging visits to P. cuspidata were not recorded in context (3). Together these contexts allowed us to explore how pollinator color preference was influenced by both plant species identity and the presence of an additional plant species in the community. For each array type we alternated flower colors in a 4 × 5 grid (fig. 1). In total, each color had the same number of open flowers (ranging from 518 to 1,254 across days). See figure 1 for schematic of array types. Due to logistical limitations we collected foraging data on one context per year (see Fig. 1 for details).

Data analysis Only plant visits from pipevine swallowtails and skippers were included in our analyses. Other pollinator species (5% of total visits) were excluded because of their small sample sizes and because their behavior on flowers suggested that they could not access the pollen or nectar rewards of the flowers. Because arrays contained equal numbers of each compared color, pollinator preference could be measured as the proportion of total floral visits to the light-blue focal species. A value of 0.5 indicates no preference, and a value greater than 0.5 indicates preference for light-blue flowers.

We used GLMMs with binomial errors and a logit link function in the lme4 package for R to model the number of visits a pollinator makes to light blue flowers versus other color flowers (Hothorn et al. 2013). We included three fixed effects in our model: foraging context (DvD,CvD DvDwC; see fig 1;), pollinator type (pipevine swallowtail or skipper) and flower color type (light-red, dark-red, dark-blue). We included pollinator individual as well as date of data collection as random effects in the model. We assessed the flexibility of pollinator color preference with a model including all two-way interactions and the three-way interaction between our three fixed effects. We determined that a model including the three-way interaction of the three main effects was the best-fit model through a likelihood ratio test. To understand the specific foraging-context by color-type by pollinator-type interactions causing this significant three-way interaction, we ran pairwise post-hoc tests using the glht() function in the multcomp package for R (Hothorn et al. 2013). We did not adjust for multiple tests because our contrasts were targeted to test specific a priori hypotheses. Implementing a correction does not change the interpretation of our results. Our contrasts were targeted to assess 1) significant differences in color preference within a given color-type across all three foraging contexts for a given pollinator and 2) differences in color preference between the two pollinator types in a particular foraging context (see tables 1 & 2 for pairwise comparisons).

**Table 1.**
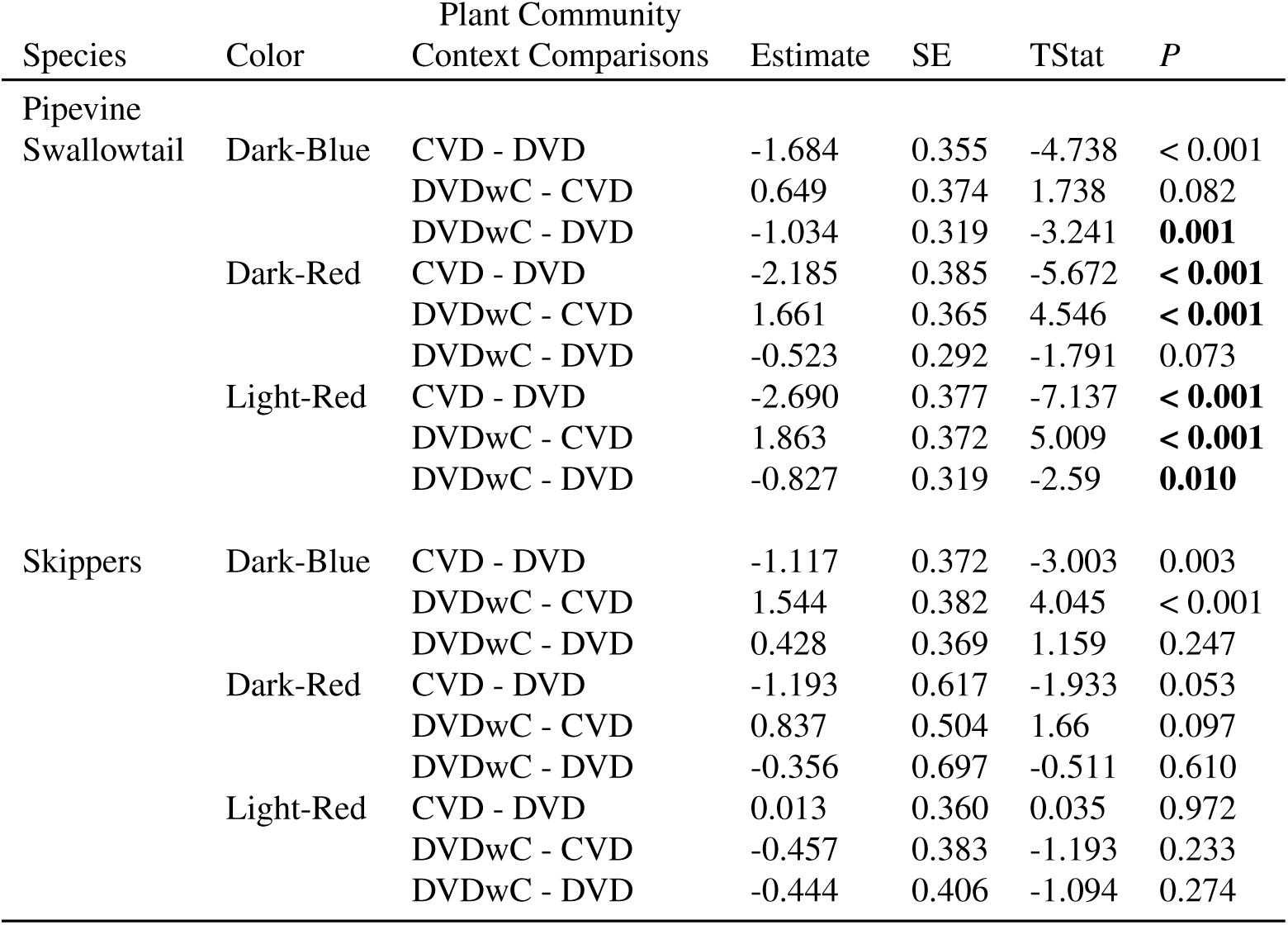
Results from post-hoc pairwise comparisons based on generalized linear mixed-effects models with binomial errors. Comparisons explore how species context influences color preference within each pollinator group. Bolded text in columns indicate significant differences in color preferences between contexts for a given pollinator group. See Figure 2 for details on which color was preferred in each context.

**Table 2.** Pollinator Comparisons: Pipevine swallowtail Skipper. Results from post-hoc pairwise comparisons based on generalized linear mixed-effects models with binomial errors. Comparisons explore how the two pollinator groups differ in their species context dependent color preference. Bolded text indicates significant differences in color preferences between swallowtail and skippers within a given context. See Figure 3 for more details about which color was preferred in each context.

| Color | Context | estimate | SE | TStat | P |
| --- | --- | --- | --- | --- | --- |
| Dark-Blue | CvD | -1.035 | 0.338 | -3.064 | 0.002 |
|  | DvD | -0.468 | 0.259 | -1.806 | 0.071 |
|  | DvDwC | -1.93 | 0.335 | -5.764 | < 0.001 |
| Dark-Red | CvD | -2.805 | 0.384 | -7.302 | < 0.001 |
|  | DvD | -1.813 | 0.564 | -3.214 | 0.001 |
|  | DvDwC | -1.98 | 0.437 | -4.528 | < 0.001 |
| Light-Red | CvD | -2.384 | 0.327 | -7.292 | < 0.001 |
|  | DvD | 0.319 | 0.308 | 1.037 | 0.300 |
|  | DvDwC | -0.064 | 0.305 | -0.209 | 0.834 |

Over three years we observed 656 pipevine swallowtail butterflies and 723 skipper butterflies foraging on our experimental arrays. The model that best predicted color preference of the two butterflies was the full model that included a three-way interaction between the fixed effects (foraging context, pollinator type, and flower color type). This model revealed a highly significant three-way interaction (see tables 1 & 2 for pairwise comparisons). For the purpose of this study we were interested in understanding how color preference and foraging context interact to shape flexibility in pollinator color preference. Furthermore we wanted to know if the two main pollinators of Phlox show different or similar color preference within each foraging context. As such, we report post hoc tests relevant for answering those specific questions below.

##### Context Dependent Preference

Pipevine swallowtails showed significant flexibility in color preference depending on foraging context and skippers showed little to no flexibility of color preference across the different foraging contexts in our study (Fig. 2). For pipevine swallowtail, the strength and direction of preference is significantly different between contexts within each of the color-type arrays (Fig. 2, Table 1). For example, in the light-red arrays, pipevine swallowtails have a strong preference for light-blue flowers when the two flower colors are the same species (DvD), no color preference when P. cuspidata is present in the array (DvDwC), and preference for light-red color when the two flower colors are different species (DvC). We observed qualitatively similar patterns of color preference flexibility for the dark-red and dark-blue color arrays as well (see Fig. 2 and Table 1 for contrast results).

**Fig. 2.**
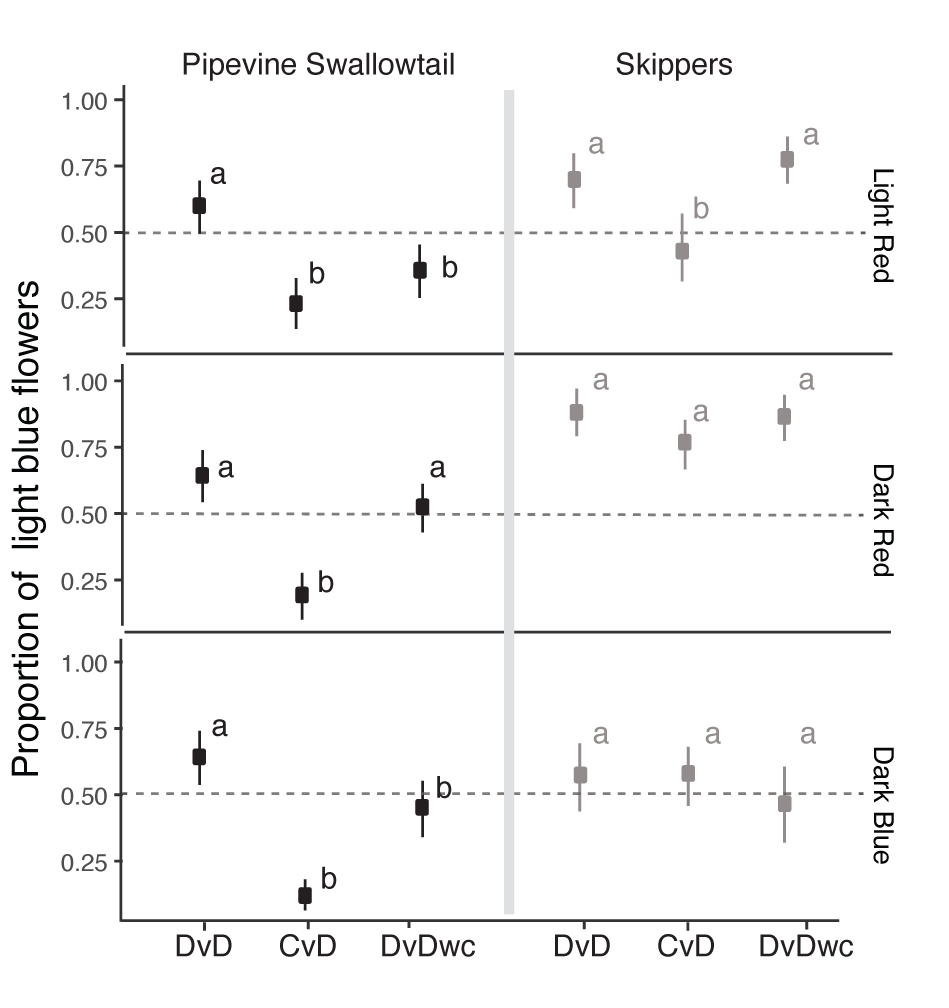
Context dependent flower color preferences vary by pollinator species. Mean proportion of visits to light blue flowers vs other color flowers across the three contexts (see figure 1 for full description of contexts) for pipevine swallowtails (in black) and skipper butterflies (in grey). 95% bootstrap CI are plotted around the mean. Letters indicate significant differences in color preferences between the three contexts for each butterfly group and color-type. Model results from the contrasts comparing the preference across contexts are displayed in table 1.

In contrast, skippers do not exhibit significant flexibility in color preference and are generally consistent with color preference regardless of the foraging context (Fig 2, Table 1). In the light-red arrays, skippers display no color preference in all three contexts. In the dark-red arrays, skippers exhibit a strong preference for light-blue flowers in all three foraging contexts. Skippers in the dark-blue arrays exhibit preference for light-blue flowers in two of the three contexts and weak to no preference when light-blue P. cuspidata is paired with dark-blue P. drummondii (DvC). *Pollinator contrasts:* The pipevine swallowtail and skippers, vary across foraging contexts in whether or not they show similar color preference (Fig. 3, Table 2). Skippers and pipevine swallowtails show similarly strong preference for light-blue P. drummondii flowers when paired with the other colors of P. drummondii (DvD). But, when P. cuspidata is paired with P. drummondii (DvC), the two butterflies show significantly different color preferences for all three color-type arrays. Similarly, in both the dark-blue and dark-red color arrays, the two pollinators have significantly different color preference for P. drummondii flower color even if P. cuspidata is just present in the background (DvDwC).

**Fig. 3.**
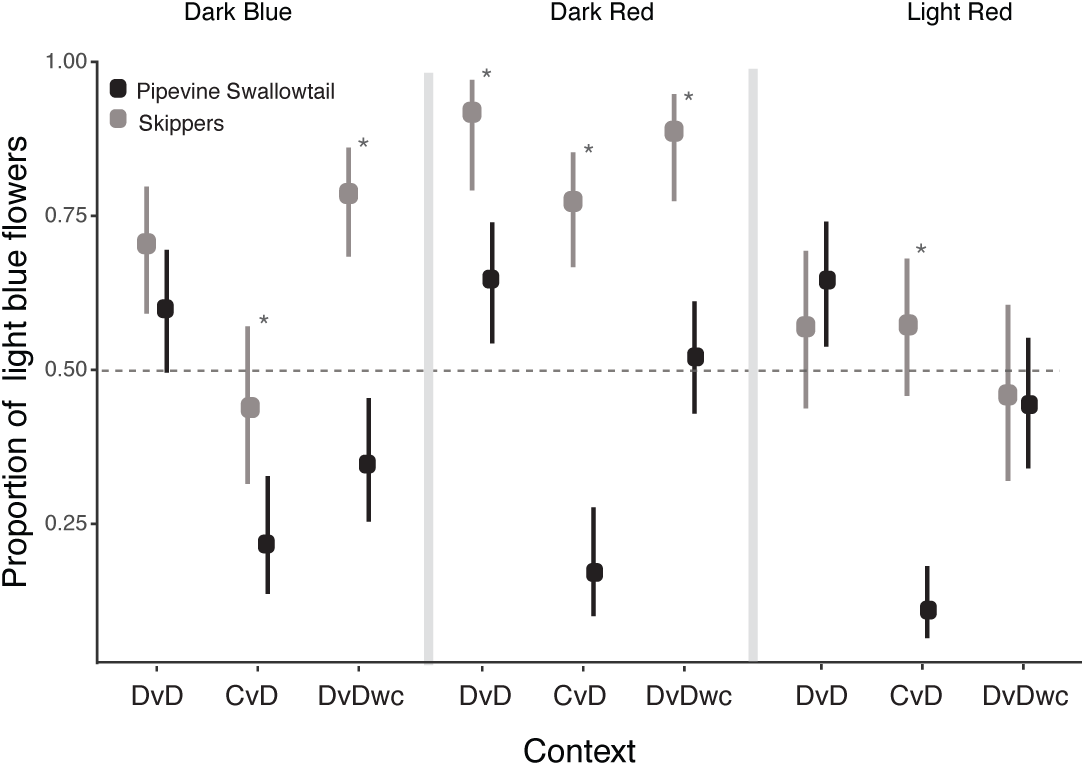
Butterfly groups differ in flower color preference across contexts. Mean proportion of visits to light-blue flowers vs other color flowers across the three contexts (see figure 1 for full description of contexts) for pipevine swallowtails (in black) and skipper butterflies (in grey). 95% bootstrap CI are plotted around the mean. Asterisks indicate significant differences in color preference between the two pollinator groups within a given context. Model results from the contrasts comparing the two butterfly groups are displayed in Table 2.

## Discussion

We found that two groups of generalist pollinators, pipevine swallowtails and skippers, vary in the consistency of their color preference while foraging in a natural field experiment. This variation in flexibility across the pollinators means that pollinator-driven selection on flower color is inconsistent across foraging contexts. Our results are based on observing wild butterflies, which have unknown foraging experience, foraging on arrays of native plants in their natural habitat. This experiment was performed using two Phlox wildflower species that depend on these pollinators for as much as 95% of their pollination visits, suggesting this behavioral variability has important implications for plant evolution. Pipevine swallowtails showed context-dependent color preference such that the strength and direction of their color preference depends on both the species identities of plants that differ in color and the presence or absence of other plant species in the area. For example, we found that these butterflies show preference for light-blue over dark-red under one foraging context, no preference in another context, and preference for dark-red over light-blue in the third context. We found similar inconsistencies in the direction and strength of preference when pipevine swallowtails choose between light-blue and light-red as well as light-blue and dark-blue flowers. These results suggest that pipevine swallowtails could use other traits and environmental signals in addition to color to make foraging decisions and are therefore flexible in their color preference under naturally variable conditions.

In contrast, the skipper butterflies did not show context dependent color preference. They displayed consistent color preferences regardless of which plant species were present or which species varied in color. Across all three color comparisons we found that skippers displayed similar strength and direction of preference regardless of the species being compared or the species present in the array. This strong preference consistency suggests that skippers use color as an important foraging cue and are relatively inflexible in their preference for particular colors.

Previous studies in this system demonstrated that flower color variation across the range of P. drummondii is maintained by pollinator-mediated selection (Hopkins and Rausher 2012; Hopkins and Rausher 2014). Dark-red flower color is favored in populations sympatric with P. cuspidata because pollinator behavior decreases costly hybridization between the two species when they have different flower colors (Hopkins and Rausher 2012). Additionally, pollinator behavior in allopatry favors light-blue flower color and thus maintains the ancestral phenotype in western P. drummondii populations (Hopkins and Rausher 2014). In this system, understanding the flexibility of pollinator behavior across plant species and community contexts is important to determine the stability of selection on flower color across geographic space and time. Our study suggests that more research is needed to understand whether flexible color preference in pipevine swallowtails leads to spatially or temporally varying selection on flower color. For example, do pipevine swallowtails in the sympatric range actually discriminate against light-blue P. drummondii plants because P. cuspidata is in the community? While P. drummondii and P. cuspidata have similar light-blue flowers, the flowers differ both in size (P. cuspidata has smaller flowers) as well as nectar amount (P. cuspidata has lower nectar volume and sugar concentration compared to P. drummondii) (Hopkins, unpubl. data). Variation in traits other than flower color could lead to context dependent preferences in pipevine swallowtails. This would suggest an additional mechanism through which pollinator-mediated selection acts on flower color.

Much of what we have learned about color preference in butterflies comes from lab studies, often explored through the use of artificial flowers (Kelber and Pfaff 1997; Weiss 1997; Kinoshita et al. 1999; Weiss and Papaj 2003; Blackiston et al. 2011). Therefore, despite the wealth of information we have about butterfly color preference, we know little about how these behaviors translate to natural systems. From these lab studies it is evident that many butterflies display innate color preferences as well as learned associations between colors and nectar rewards. For example, pipevine swallowtails have an innate preference for blue flowers over yellow in the lab (Weiss 1997). In our study we found that while pipevine swallowtails preferred blue flowers of one species (P. drummondii), but were strongly deterred by the blue flowers of another species (P. cuspidata), and ultimately displayed context dependent preference for blue flowers. These results suggest that our understanding of pipevine swallowtail flower color preference from the lab does not necessarily translate to behavior we observed in the field and that these butterflies are likely using other cues in addition to color (such as shape or scent) to guide their foraging preferences. In contrast we found that skippers more consistently base their foraging decisions on flower color, as color preference did not appear to be influenced by foraging context. The few field studies that examine butterfly color preference suggest that contextdependent color preference may be common (Clements 1923; Pohl et al. 2011) making the case for future studies that explore color preference in natural contexts.

Pollinator preference for particular floral traits exerts selective pressures on plants. It is therefore of primary interest to understand the extent to which co-occurring pollinators exert either similar or disparate selective forces on plants. In this study we found that in some foraging contexts pipevine swallowtails and skippers exhibit the same color preferences and in other contexts we found that the two pollinators display disparate color preferences. In other words, whether or not the two pollinator groups impose concordant selection on flower color depends on the species being compared and the background community of plant species. In specialized plant pollinator interactions, conclusions as to how a pollinator acts as an agent of selection on specific floral traits can be relatively straightforward (i.e. Muchhala and Thomson 2009). However, most plants are visited by multiple pollinator species and the strength and direction of selection multiple pollinators impose on floral traits is rarely assessed. Furthermore, the composition of the pollinator visitors can vary both spatially (Gomez et al. 2010) and temporally (CaraDonna et al. 2017), further complicating our understanding of how pollinator variation drives patterns of selection on floral traits. Understanding the flexibility of pollinator behavior across plant species and community contexts is crucial for determining the stability of selection on flower color across geographic space and time.

The two butterflies groups in our study showed different degrees of flexibility in their color preference. These differences in color preference can be due to a number of factors including differences in visual systems and/or differential learning abilities. Unlike most other groups of pollinators, visual pigments of butterfly eyes are varied across families and even species. This means that butterfly individuals of different species can both collect and perceive spectral information in different ways. Not surprisingly, this variation in color perception can lead to differences in innate color preferences, the ability to learn new colors associated with rewards, as well as the degree to which color preference will be context dependent and influenced by the environment (Briscoe 2008; Blackiston et al. 2011). While some butterflies have red visual receptors, it appears that skippers do not, likely leading to passive discrimination against red colored flowers (Briscoe and Chittka 2001). Currently there are no studies examining the pipevine swallowtail visual system, but closely related species exhibit exceptional long-wavelength visual abilities (Arikawa 2003; Takemura et al. 2005). Future studies that link flexibility in preference to variation in visual systems will add an invaluable mechanistic understanding of how butterflies can impact selection on plant traits.

The frequency and pattern of pollinator foraging can have a direct impact on plant reproductive success and the evolution of plant traits and yet we know little about mechanisms behind pollinator behavior. Many lab studies suggest that butterflies display innate and learned color preferences and that these preferences can be flexible, but our study is one of few that explores the flexibility of color preference in the field. Therefore it remains unclear how results from lab studies translate to behavior in natural systems. Our study reveals that two butterfly groups that provide the majority of pollination visitation to two native wildflowers display different flexibility in color preference and, in the case of the pipevine swallowtail, behave in ways that might be difficult to predict from lab studies. This study enhances our understanding of if and how pollinators display flexible foraging preferences in the wild. Future studies that combine descriptions of visual systems with critical behavioral assays in the lab and in natural environments will allow us to understand the prevalence and mechanisms underlying flexibility in pollinator foraging behavior.

## ACKNOWLEDGEMENTS

We thank the Brackenridge field laboratory for field experimental support. Thanks to University of Texas at Austin Greenhouse manager Shane Merrell and the staff of Harvard’s Arnold Arboretum for assistance with plant maintenance. RH received funding from the William F. Milton Fund through Harvard University. SG was funded by a Category B Erasmus Mundus scholarship. CMS was supported by National Defense Science & Engineering Graduate Fellowship (NDSEG). Thanks also to D. Papaj for his comments on the manuscript.

## References

Aldridge, G. & Campbell, D.R. (2007) Variation in pollinator preference between two Ipomopsis contact sites that differ in hybridization rate. Evolution, 61, 99–110.

Arikawa, K. (2003) Spectral organization of the eye of a butterfly, Papilio. Journal of comparative physiology A, Neuroethology, sensory, neural, and behavioral physiology, 189, 791–800.

Blackiston, D., Briscoe, A.D. & Weiss, M.R. (2011) Color vision and learning in the monarch butterfly, Danaus plex-DRAFT REFERENCES ippus (Nymphalidae). The Journal of experimental biology, 214, 509–520.

Briscoe, A.D. (2008) Reconstructing the ancestral butterfly eye: focus on the opsins. The Journal of experimental biology, 211, 1805–1813.

Briscoe, A.D. & Chittka, L. (2001) The evolution of color vision in insects. Annual Review Of Entomology, 46, 471–510.

Brosi, B.J. & Briggs, H.M. (2013) Single pollinator species losses reduce floral fidelity and plant reproductive function. Proceedings of the National Academy of Sciences, 110, 13044–13048.

Byers, K.J.R.P., Bradshaw, H.D. & Riffell, J.A. (2014) Three floral volatiles contribute to differential pollinator attraction in monkeyflowers (Mimulus). The Journal of experimental biology, 217, 614–623.

CaraDonna, P.J., Petry, W.K., Brennan, R.M., Cunningham, J.L., Bronstein, J.L., Waser, N.M. & Sanders, N.J. (2017) Interaction rewiring and the rapid turnover of plant-pollinator networks. Ecol Lett, 20, 385–394.

Clements, F.E. (1923) Experimental Pollination An Outline Of The Ecology Of Flower And Insects.

Dötterl, S., Glück, U., Jürgens, A., Woodring, J. & Aas, G. (2014) Floral Reward, Advertisement and Attractiveness to Honey Bees in Dioecious Salix caprea. PLoS ONE, 9, e93421.

Fenster, C.B., Reynolds, R.J.,Williams, C.W., Makowsky, R. & Dudash, M.R. (2015) Quantifying hummingbird preference for floral trait combinations: The role of selection on trait interactions in the evolution of pollination syndromes. Evolution, 69, 1113–1127.

Fornoff, F., Klein, A.M., Hartig, F., Benadi, G., Venjakob, C., Schaefer, H.M. & Ebeling, A. (2016) Functional flower traits and their diversity drive pollinator visitation. Oikos.

Giurfa, M., N ez, J., Chittka, L. & Menzel, R. (1995) Colour preferences of flower-naive honeybees. Journal of Comparative Physiology A, 177.

Gomez, J.M., Abdelaziz, M., Lorite, J., JesÚs MuÃ±oz-Pajares, A. & Perfectti, F. (2010) Changes in pollinator fauna cause spatial variation in pollen limitation. Journal Of Ecology, 98, 1243–1252.

Goulson, D., Cruise, J.L., Sparrow, K.R. & Harris, A.J. (2007) Choosing rewarding flowers; perceptual limitations and innate preferences influence decision making in bumblebees and honeybees. Behavioral Ecology And Sociobiology, 61, 1523–1529.

Gumbert, A. (2000) Color choices by bumble bees (Bombus terrestris): innate preferences and generalization after learning. Behavioral Ecology And Sociobiology, 48, 36–43.

Hersch, E.I. & Roy, B.A. (2007) Context-dependent pollinator behavior: an explanation for patterns of hybridization among three species of indian paintbrush. Evolution, 61, 111–124.

Hollis, K. & Guillette, L. (2011) Associative learning in insects: Evolutionary models, mushroom bodies, and a neuroscientific conundrum. Comparative Cognition & Behavior Reviews, 6, 24–45.

Hopkins, R. & Rausher, M.D. (2012) Pollinator-Mediated Selection on Flower Color Allele Drives Reinforcement. Science, 335, 1090–1092.

Hopkins, R., Guerrero, R.F., Rausher, M.D. & Kirkpatrick, M. (2014) Strong reinforcing selection in a Texas wildflower. Current biology : CB, 24, 1995–1999.

Hopkins, R. & Rausher, M.D. (2011) Identification of two genes causing reinforcement in the Texas wildflower Phlox drummondii. NATURE, 469, 411–414.

Hopkins, R. & Rausher, M.D. (2014) The Cost of Reinforcement: Selection on Flower Color in Allopatric Populations of Phlox drummondii*. The American Naturalist, 183, 693–710.

Hothorn, T., Bretz, F., Westfall, P., Heiberger, R.M. & Schuetzenmeister, A. (2013) Package “multcomp”. 2013.

Ings, T.C. & Chittka, L. (2009) Predator crypsis enhances behaviourally mediated indirect effects on plants by altering bumblebee foraging preferences. Proceedings Biological sciences / The Royal Society, 276, 2031–2036.

Johnsen, S., Kelber, A. & Warrant, E. (2006) Crepuscular and nocturnal illumination and its effects on color perception by the nocturnal hawkmoth Deilephila elpenor. The Journal of experimental biology, 209, 789–800.

Kandori, I., Yamaki, T., Okuyama, S.I., Sakamoto, N. & Yokoi, T. (2009) Interspecific and intersexual learning rate differences in four butterfly species. The Journal of experimental biology, 212, 3810–3816.

Kelber, A. & Pfaff, M. (1997) Spontaneous and learned preferences for visual features in a diurnal hawkmoth. Israel Journal of Plant Sciences, 45, 235–245.

Kelber, A. & Pfaff, M. (1999) True Colour Vision in the Orchard Butterfly, Papilio aegeus. Naturwissenschaften, 86, 221–224.

Kinoshita, M., Shimada, N. & Arikawa, K. (1999) Colour vision of the foraging swallowtail butterfly papilio xuthus. The Journal of experimental biology, 202 (Pt 2), 95–102.

Knauer, A.C. & Schiestl, F.P. (2014) Bees use honest floral signals as indicators of reward when visiting flowers. Ecol Lett.

Leonard, A.S. & Masek, P. (2014) Multisensory integration of colors and scents: insights from bees and flowers. Journal of Comparative Physiology A, 200, 463–474.

Levin, D.A. & Berube, D.E. (1972) PHLOX AND COLIAS: THE EFFICIENCY OF A POLLINATION SYSTEM. Evolution, 26, 242–250.

Muchhala, N. (2003) Exploring the Boundary between Pollination Syndromes: Bats and Hummingbirds as Pollinators of Burmeistera cyclostigmata and B. tenuiflora (Campanulaceae). Oecologia, 134, 373–380.

Muchhala, N. & Thomson, J.D. (2009) Going to great lengths: selection for long corolla tubes in an extremely specialized bat–flower mutualism. Proceedings Of The Royal Society B-Biological Sciences, 276, 2147–2152.

Ollerton, J., Alarcón, R., Waser, N.M., Price, M.V., Watts, S., Cranmer, L., Hingston, A., Peter, C.I. & Rotenberry, J. (2009) A global test of the pollination syndrome hypothesis. Annals of Botany, 103, 1471–1480.

Ômura, H. & Honda, K. (2005) Priority of color over scent during flower visitation by adult Vanessa indica butterflies. Oecologia, 142, 588–596.

Osorio, D. & Vorobyev, M. (2008) A review of the evolution of animal colour vision and visual communication signals. Vision research, 48, 2042–2051.

Pohl, N.B., VanWyk, J. & Campbell, D.R. (2011) Butterflies show flower colour preferences but not constancy in foraging at four plant species. Ecological Entomology, 36, 290–300.

Raguso, R.A. (2008) Wake Up and Smell the Roses: The Ecology and Evolution of Floral Scent. Annual Review Of Ecology Evolution And Systematics, 39, 549–569.

Raine, N.E. & Chittka, L. (2007) Flower constancy and memory dynamics in bumblebees (Hymenoptera: Apidae: Bombus). ENTOMOLOGIA GENERALIS, 29, 179–199.

Rosas-Guerrero, V., Aguilar, R., Martén-Rodríguez, S., Ashworth, L., Lopezaraiza-Mikel, M., Bastida, J.M. & Quesada, M. (2014) A quantitative review of pollination syndromes: do floral traits predict effective pollinators? Ecol Lett, 17, 388–400.

Russell, A.L., Newman, C.R. & Papaj, D.R. (2016) White flowers finish last: Pollen-foraging bumble bees show biased learning in a floral color polymorphism. Evolutionary Ecology, pp. 1–20.

Schemske, D.W. & Bradshaw, H.D. (1999) Pollinator preference and the evolution of floral traits in monkeyflowers (Mimulus). Proceedings of the National Academy of Sciences, 96, 11910–11915.

Schiestl, F.P. & Johnson, S.D. (2013) Pollinator-mediated evolution of floral signals. Trends in Ecology & Evolution, 28, 307–315.

Shrestha, M., Dyer, A.G., Boyd-Gerny, S., Wong, B.B.M. & Burd, M. (2013) Shades of red: bird-pollinated flowers target the specific colour discrimination abilities of avian vision. New Phytologist, 198, 301–310.

Smith, S.D., Ané, C. & Baum, D.A. (2008) The role of pollinator shifts in the floral diversification of Iochroma (Solanaceae). Evolution, 62, 793–806.

Stavenga, D.G. & Arikawa, K. (2006) Evolution of color and vision of butterflies. Arthropod structure & development, 35, 307–318.

Takemura, S.Y., Kinoshita, M. & Arikawa, K. (2005) Photoreceptor projection reveals heterogeneity of lamina cartridges in the visual system of the Japanese yellow swallowtail butterfly, Papilio xuthus. The Journal of comparative neurology, 483, 341–350.

Thairu, M.W. & Brunet, J. (2015) The role of pollinators in maintaining variation in flower colour in the Rocky Mountain columbine, Aquilegia coerulea. Annals of Botany, 115, 971–979.

von Frisch, E., Lindauer, M. & Daumer, K. (1914) On the perception of polarized light through bees’ eyes. Experientia, 16, 289–301.

Waser, N.M., Chittka, L., Price, M., Williams & Ollerton, J. (1996) Generalization in pollination systems, and why it matters. Ecology, 77, 1043–1060.

Weiss, M.R. (1997) Innate colour preferences and flexible colour learning in the pipevine swallowtail. Animal Behaviour, 53, 1043–1052.

Weiss, M.R. & Papaj, D.R. (2003) Colour learning in two behavioural contexts: how much can a butterfly keep in mind? Animal Behaviour, 65, 425–434.

Yoshida, M., Itoh, Y., Ômura, H., Arikawa, K. & Kinoshita, M. (2015) Plant scents modify innate colour preference in foraging swallowtail butterflies. Biology Letters, 11.

